# Use of deep learning algorithms for segmentation of microelectrode arrays electrograms in the study of changes in myocardial bioelectrical activity

**DOI:** 10.1101/2024.11.19.624266

**Authors:** Elena Kotikhina, Denis Karchkov, Viktor Moskalenko, Anton Rybkin, Ruslan Smirnov, Grigory Osipov, Lev Smirnov

## Abstract

The using of modern electrophysiological methods during scientific research implies processing of huge data sets which becomes a significant problem. Progress in the development of equipment for registration of multiple local field potentials from the tissue surface contributes to the study of spatio-temporal characteristics of myocardial bioelectrical activity at a more detailed level. But it also raises the need for automation of data analysis, which can be accomplished by algorithms based on machine learning. New scientific approaches can contribute to the discovery of new facts about substances widely used but with under-researched efficacy. The paper presents the results of epicardial mapping by flexible microelectrode arrays of myocardium of isolated rat heart perfused with a solution containing L-carnitine, nutritional supplement that may be recommended for people suffering from cardiovascular disease. Electrograms from microelectrode arrays were analyzed using a neural network model based on U-Net architecture adapted for segmentation of one-dimensional signal. It was shown that the presence of L-carnitine in the perfusion solution caused a decrease in heart rate, myocardial excitation conduction velocity, coronary blood flow intensity of the isolated rat heart and suppressed physiological responses of the heart to adrenaline stimulation.

## Introduction

The problem of analyzing the large amount of data recorded during electrophysiological experiments has existed since the first studies that were carried out using a string galvanometer. T. Lewis and M.A. Rothschild in 1915 [1], arranging multiple electrodes on the surface of the dog heart (Fig. 1, A), recorded electrograms to study the myocardial bioelectrical activity (MBA) (Fig. 1, B). Modern microelectrode arrays (MEA) (Fig. 1, C) can include tens and tens of thousands of electrodes that simultaneously record bioelectrical potentials (Fig.1, D), resulting in great amount of data [2].

**Figure 1.**
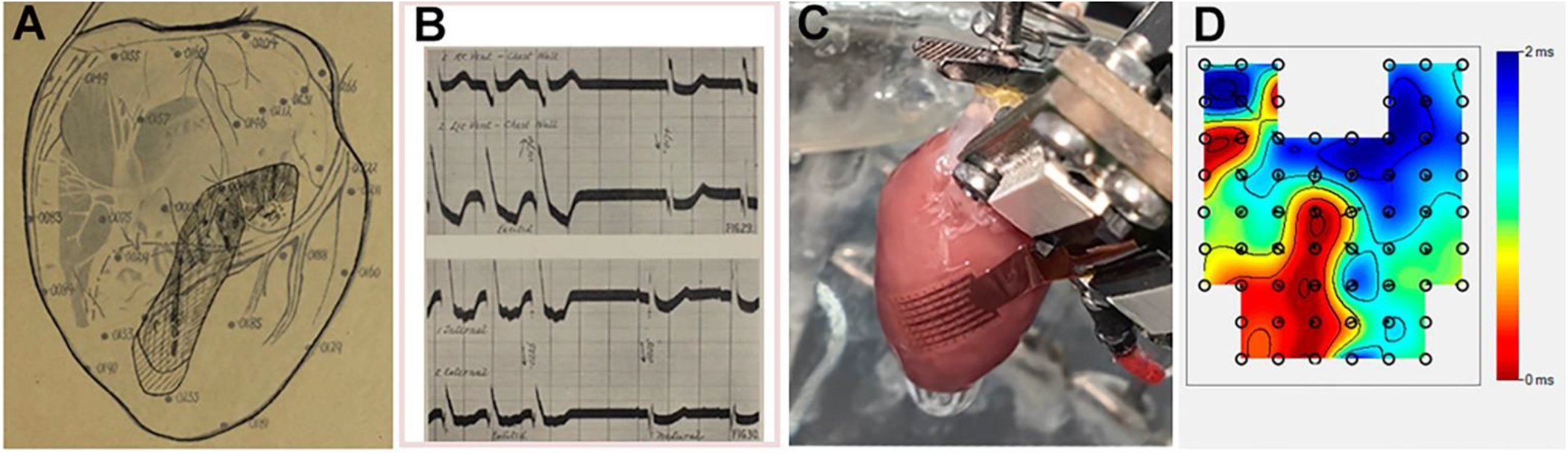
Multielectrode mapping of MBA: A - scheme of arrangement of multiple electrodes on the dog heart; B - resulting electrograms [1]; C - MBA registration of the isolated rat heart by FlexMEA72 (Multichannel systems, Germany); D - isochronal activation map of MBA obtained by this method

Algorithms based on deep learning (DL) have been used to solve problems in the fields of cardiology and neurobiology, where membrane and field potential (FP) parameters are investigated [3-10]. Cardiac data can be represented as a large array of tissue excitation moments recorded from different points from the epicardium, endocardium, or myocardial slice, enabling investigation of spatio-temporal changes in MBA, identification of ectopic foci and zones of electrical non-conductivity of cardiac tissue, and characterization of local field potentials (LFPs) [6, 7]. In experimental conditions it becomes possible to simulate physical, biochemical, pharmacological and other types of external influence on the heart. Variability of parameters of recorded bioelectrical potentials (shape, amplitude, action potentials frequency etc.), as well as the presence of electrical noise and artifacts during the experiment force the researcher to constantly adapt the settings of signal processing. Automation of electrogram signal segmentation will significantly facilitate and accelerate the process of analyzing experimental data.

One of the most effective methods of studying myocardial bioelectrical characteristics is the method of using MEA, which is characterized by high spatial resolution of the extracellular potentials recording. The method of multi-electrode mapping allows analyzing spatial and temporal parameters of MBA, in particular, the rate of conduction of electrical excitation, significant changes in which can be considered as predictors of life-threatening arrhythmias [11-22].

Flexible MEAs are used to record MBA from the heart surface [12, 23, 24]. MEA electrodes simultaneously record electrograms, representing time dependence of voltage. The microelectrodes record LFPs - a signals which contain both action potentials and other membrane potential-derived fluctuations in a small extracellular volume [25, 26]. Activation time (AT) - the maximum steepness of the LFP decline (dV/dt), corresponds to the occurrence of an action potential on cell membrane in the region of recording electrode positioning [27, 28]. Analyzing the frequency of ATs occurrence at a single electrode or the sequence of occurrence at multiple electrodes within the MEA makes it possible to estimate such parameters as pacemaker activity and myocardial electrical conductivity.

There is a large number of software for operating with data obtained by the MEA method, providing conversion, visualization of the electrical signal, filtering of electrical noise, and primary processing [29-35]. The development of a universal algorithm for automating the analysis of MEA electrograms would not only help to accelerate the final results of the research, but also simplify their comparative evaluation and increase their provability. Such an algorithm can be based on the application of DL methods. And experimental data acquisition can provide a large and diverse sample of bioelectrical potential parameters for training neural networks.

The paper presents the results obtained by epicardial mapping of MBA of the isolated rat heart by flexible MEA. For electrograms segmentation, a neural network model based on the U-Net architecture modified for processing one-dimensional signals was applied. The U-Net architecture involves convolutional layers for feature extraction and a decoder for accurate reconstruction of spatial information. The activity of the heart provides blood circulation in the organism to maintain its vital activity and requires large energy consumption. According to various estimations, the amount of adenosine triphosphate (ATP) produced and utilized in the heart per day exceeds the mass of the organ by 15-100 times [36, 37]. Among all cell types, cardiomyocytes are the richest in mitochondrial density, up to one-third of the cell volume [38, 39]. Such processes specific for cardiomyocytes as the work of actin-myosin complex and transmembrane ion currents against its concentration gradient by ATP-dependent pumps to maintain cell membrane potential and to store calcium ions in the sarcoplasmic reticulum [40] occur with energy consumption.

Under normal conditions, more than 95% of ATP, the main and universal carrier of chemical energy in the cell, in the adult mammalian heart is synthesized by oxidative phosphorylation of fatty acids in the mitochondria, and 5% by glycolysis in the cytoplasm [41-45]. The transfer of long-chain fatty acids into the mitochondrial matrix for energy synthesis in the form of ATP in the process of β-oxidation is carried out with the participation of L-carnitine (LC) [46]. LC (beta-hydroxy-gamma-N-trimethyl-aminobutyric acid) is synthesized in the human body from the amino acids lysine and methionine predominantly in the liver and kidneys, but it is most abundant in skeletal muscle and myocardium, where it is transported from blood plasma [47-51]. The bioavailability of LA depends on its content in food and ranges from 54 to 87%. On average, a person with a dietary intake of red meat, the richest source of LC, obtains 1-2.4 mg (6-15 µMol) of LC per 1 kg body weight per day [52]. Pharmacologic supplemental doses of LC administered orally in the form of tablets or solutions are less bioavailable (5-18%). Intravenous administration can increase the plasma concentration of LC to a greater degree, but after 12-24 hours it returns to a norm of 40-60 μMol/L through kidney function [53]. Some studies suggest that long-term administration (6 months), especially in combination with carbohydrates, can increase the content of total LC in skeletal muscles [54, 55]. In the body of an adult healthy person, there is no functional deficiency of LC accompanied by abnormal clinical manifestations with the possibility of correction by prescribing additional doses of LC [56].

Due to its important role in the process of ATP synthesis from fatty acids, LC (as well as its derivatives, e.g. propionyl-LC) has come to be considered as a potentially effective supplement for improving both skeletal muscle and myocardial performance [57-59]. Hypothetically, an increase in the content of LC and its derivatives in the cell should increase the intensity of reactions of energy metabolism, and thus the contractile activity and endurance of myocardium.

The positive effect of supplementation with additional doses of LC on myocardial function in healthy organism and in the presence of cardiovascular diseases (CVD) may be related to its antioxidant [60, 61] and anti-inflammatory effects [62]. LC provides transport of long-chain fatty acids into mitochondria, excessive accumulation of which in the cytosol of cardiomyocytes can lead to diastolic dysfunction and prolongation of the QT interval on the electrocardiogram due to suppression of repolarizing K^+^-currents [63]. Additional LC supplementation can maintain an optimal ratio of carnitine and its esters in the heart, thereby reducing the degree of aggressive effects of the latter on cell membranes [64, 65]. Increasing LC availability reduces the intramitochondrial acetyl-CoA/CoA ratio and thereby increases glucose utilization while suppressing fatty acid consumption as an energy substrate in the heart [66].

To date, the results of studies using meta-analysis on large groups of patients suggest that supplementation with LC may be effective in the treatment and prevention of cardiovascular pathologies and risk factors for their development [67]. LC treatment of chronic heart failure was effective, showing improvements in clinical symptoms and cardiac function, as well as a decrease in serum levels of brain natriuretic peptide and its N-terminal propeptide, markers of myocardial dysfunction [68]. The beneficial effects of LC supplementation on the cardiovascular system have been shown by reducing oxidative stress and inflammation indicators in patients with coronary heart disease [69]. LC and its derivatives have favorable effects on vascular function by influencing endothelial state and nitric oxide release, in particular during and after ischemia [70-73].

During LC supplementation, it should be considered that it is a nutrient substrate for the gut microbiota that produces trimethylamine-N-oxide, the elevation of level of which in the body increases the risk of atherosclerosis and subsequent CVD [74-78].

Cardiac function is regulated by the autonomic nervous system and is affected by neurotransmitters and hormones. One of the most important regulators of myocardial activity is adrenaline (ADR). An increase in blood concentrations of ADR is involved in triggering the «fight or flight» response and causes an increase in heart rate (HR) and force of contractions [79]. In healthy individuals, the action of ADR under normal conditions has no adverse implications. But for a person suffering from CVD, an increase in the intensity of cardiac work provoked by emotional stress or excessive physical activity may cause aggravations [80]. Since LC can be recommended in the treatment and prevention of CVDs, including intensive exercise as a muscle endurance-enhancing active supplement [67], [81], it is important to determine how this compound acts on myocardium affected by ADR. And taking into account the complex metabolism of LC and its wide spectrum of action in the organism, it is reasonable to study *ex vivo* the effect of increase of LC concentration on MBA and intensity of coronary circulation.

The aim of the study was to investigate the effect of additional LC on changes in MBA and coronary blood flow (CBF) intensity of the isolated rat heart under the influence of ADR. The epicardial electrograms (EE) registered with MEA were analyzed using algorithms based on DL.

## Methods

The research was performed on isolated perfused hearts of Wistar rats of both sexes. The housing, care and handling of the animals were approved by the Ethical Commission of Lobachevsky University and were in accordance with the «Guide for the Care and Use of Laboratory Animals (ILAR publication, 1996, National Academy Press)», the interstate standard GOST 33216-2014 «Guide for the Housing and Care of Laboratory Animals. Rules of keeping and care of laboratory rodents and rabbits», sanitary and epidemiological rules SP 2.2.1.3218-14.

### Langendorff method

Isolated rat hearts were perfused by the Langendorff method with Krebs-Henseleit solution (NaCl 118, KCl 4.7, CaCl_2_ 2, MgSO_4_ 1.2, KH_2_PO_4_ 1.2, NaHCO_3_ 20, glucose 10 mMol) at 37°C, 100 cm water column pressure, and carbogen bubbling. CBF volume, ml per min, was analyzed by measuring the outflow solution from the perfused heart. Depending on the experimental conditions, two groups «ADR» (n=12) and «LC» (n=12) were formed. After a period of adaptation to standard perfusion (SP) conditions - perfusion without additional influences - hearts from the «LC» group were perfused with a solution containing LC (C=1 mMol) for 15 minutes, then ADR (C=2 μMol) was injected into aorta and perfusion continued for another 10 minutes. For the «ADR» group, the protocol was the same but without LC in the perfusion solution.

### MEA method

For multichannel recording of MBA, a flexible MEA (FlexMEA72, Multichannel systems, Germany) (Fig. 2, A) was placed on the epicardium of the left ventricle (Fig. 2, B) of the isolated hearts. FlexMEA72 includes 64 recording, 4 reference and 4 grounding electrodes. Thus, EE registration with sampling frequency of 20 kHz, with filtering (1-300 Hz), amplification (50-fold) and analog-to-digital conversion (16 bits) of bioelectrical signals was performed. Cardio2D software (Multichannel systems, Germany) was used for EE visualization and recording. Activation maps were generated based on the detected ATs on EEs to measure the spatio-temporal characteristics of MBA (Fig. 2, C).

**Figure 2.**
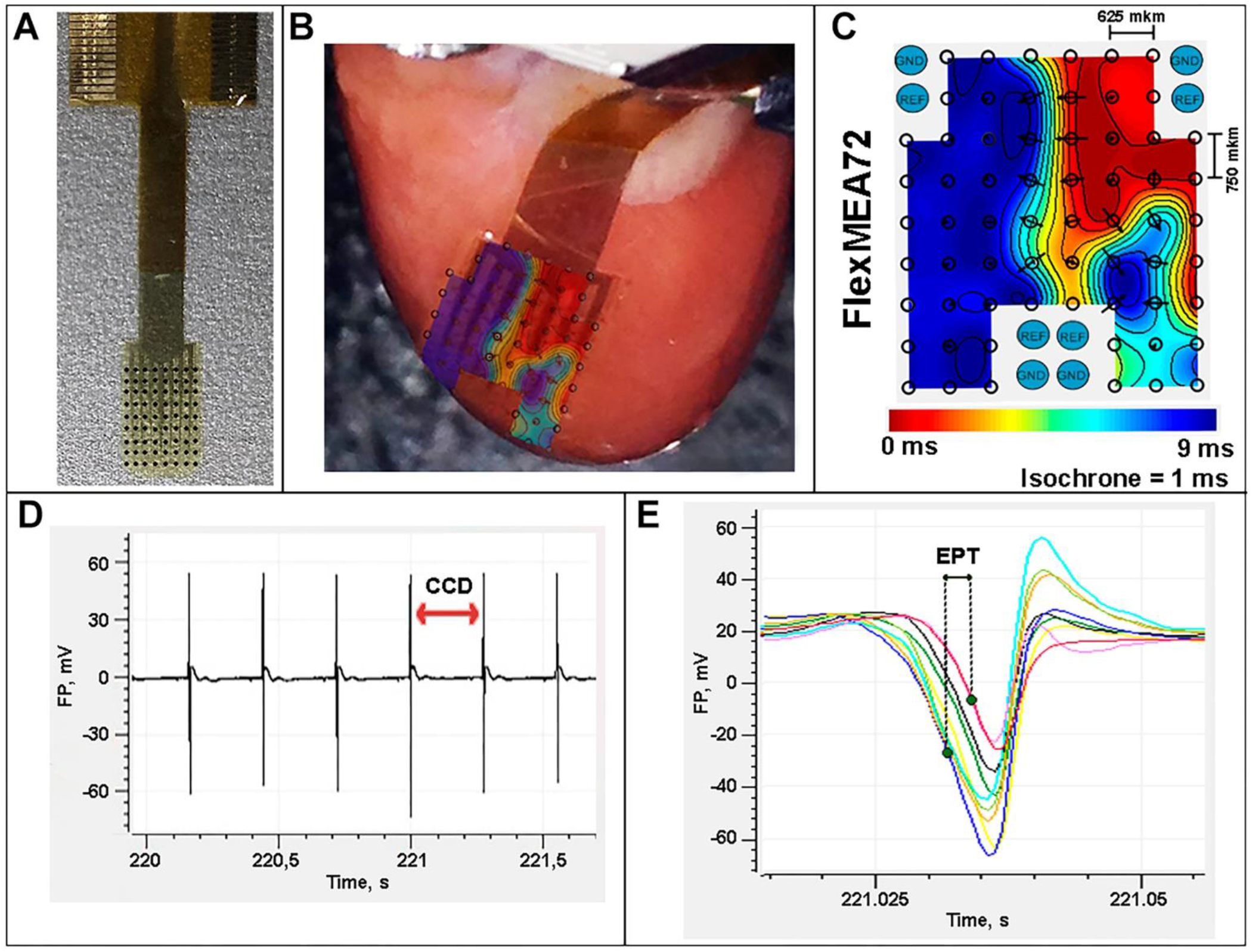
MEA mapping of MBA of the isolated rat heart: A - FlexMEA72; B - location of FlexMEA72 on the left ventricle of the isolated rat heart; C - activation map of MBA during a single heartbeat; D - sequence of complexes of LFP changes, by the time interval between ATs on which the CCD value can be determined; E - LFPs from nine recording electrodes of MEA, on the first and the last (by time of occurrence) of which ATs are labeled to determine EPT by the value of the time interval between them

The cardiac cycle duration (CCD), the time interval between the two subsequent ATs recorded by one MEA electrode, was analyzed (Fig. 2, D). The coefficient of variation of CCD (CV_CCD_) was taken as an index of HR stability:

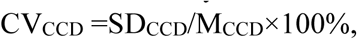

where SD_CCD_ is the standard deviation of CCD, M_CCD_ is the average value of CCD.

Excitation propagation time (EPT) was calculated as the time interval between the first and last recorded ATs at different MEA electrodes during a single heartbeat (Fig. 2, E).

CCD, CV_CCD_, and EPT were calculated using 100 values for each considered time period. Statistical data processing was performed in GraphPad Prism (GraphPad Software, USA) using two-way ANOVA (and Tukey’s test), Friedman test (and Dunn’s test) and Wilcoxon test.

### DL based algorithms for MEA data analysis

Graphical representation and primary analysis of EE were performed using Cardio2D+ software (Multichannel systems, Germany) (Fig. 3, A). To estimate the MBA parameters, it is necessary to detect ATs on the LFPs. In order to automate this procedure, a neural network model based on U-Net architecture adapted for segmentation of one-dimensional signal was developed.

**Figure 3.**
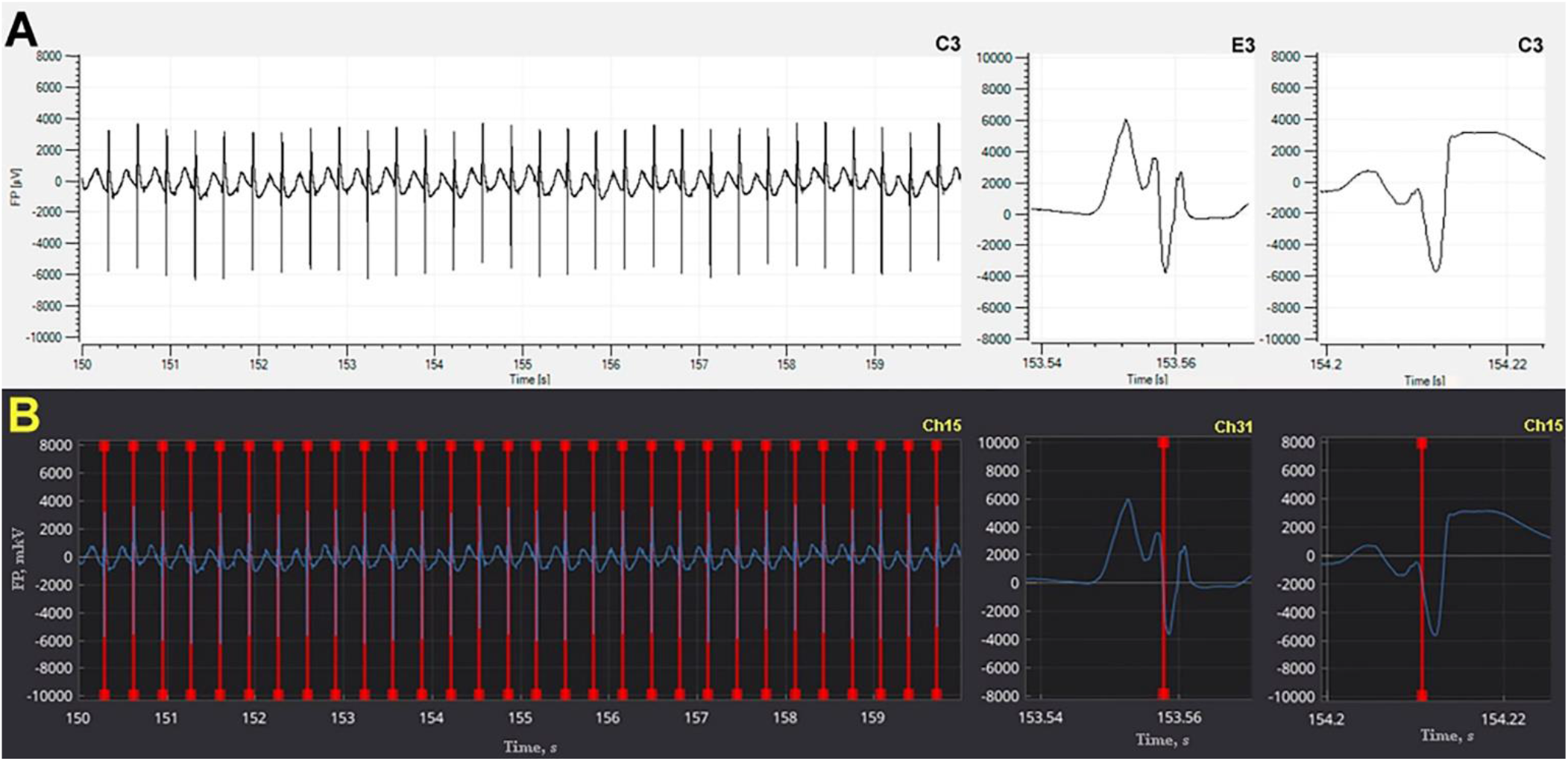
EE (dependence of field potential, FP, on time) of MBA of the isolated rat heart: A - visualization by Cardio2D+ software before labeling; B - labeling of ATs on LFP using «EGM Analyzer» software; C3, E3 – electrodes of MEA and corresponding channels Ch15, Ch31

EEs obtained in previous studies by the authors of this study ([24] Kharkovskaya et al., 2020) were used as a dataset for training, validation and testing of the model. Experimental recordings of MBA under SP conditions of the isolated rat hearts were selected, as well as recordings of hearts undergoing reversible chemically-induced ventricular fibrillation containing LFPs with high instability in amplitude, shape, and frequency of ATs occurrence. The dataset included a total of eight files (four under SP conditions and four during arrhythmia) with MBA recordings of the isolated hearts of four rats. The duration of the recordings was 10 minutes and contained EEs from 64 channels. The primary labeling obtained with Cardio2D+ software was incomplete and uncertain due to the difficult nature of the EE signal for segmentation by algorithms based on direct methods. Therefore, it was adjusted and finalized using a specially developed software «EGM Analyzer» [82, 83] (Fig. 3, B). Thus, taking into account the frequency of contractions of the isolated rat hearts, the quantity of recording channels in the MEA, some of which could register an unacceptable bioelectrical signal for processing, the training sample volume amounted to about 1,400,000 ATs.

The EE data were converted and reduced to a signal size of 2 sec (10,000 samples) and subjected to filtering and scaling. The EE recordings from each channel containing ATs were divided into a portion (12.500 samples) for further processing and a portion for resubmission to the algorithm input. A value between 0 and 2,500 samples was equally chosen as an indentation followed by sampling in 10,000 steps. Next, a binary search for labeling was performed, adding to the dataset in case of detection and in order to exclude unlabeled EE sections from the sample.

For realization of ATs localization on EEs was used Python programming language and such libraries as PyTorch (model building and training, prediction), NumPy (reading/writing and preprocessing of EEs and of labeling), SciPy (preprocessing of EEs), mlflow (storing information about experiments and controlling metrics values), ONNX, DearPyGui (graphical interface for some auxiliary programs). The source code with the realization of the described functionality is available [84].

As the architecture of the neural network model for the signal segmentation task, a U-Net type model was selected and modified. Mirror-filled padding was used to preserve the vector size during convolutions and to equalize the input and output sizes. To reduce the training and prediction time, upsample procedure was used instead of inverse convolutions in the reconstruction phase.

The neural network model did not include fully connected layers and consisted only of convolutions, pooling layers and upsample layers. The descending part of the model is represented by two consecutive convolutions and a max pooling layer, while the ascending part performs an inverse operation in which upsample is used instead of max pool. Residual links in the model architecture connect the downsample and upsample parts by concatenating the tensors. Dropout layers were applied before max pooling layers to prevent overtraining and increase the generalizability of the model [85]. During model training, adam with milestone scheduler for dynamic control of learning rate was used as an optimization method [86, 87]. The EE data for training were divided into training (six random files), validation (one random file), and test samples (1 file). Data cutting was performed as follows: a channel (MEA electrode) was randomly selected in a randomly selected file; a point was randomly selected on the EE from the selected channel; an offset relative to the selected point was randomly selected; a 2-second fragment of the bioelectrical signal was cut relative to this offset (10,000 signal samples) with segmentation corresponding to this signal. According to this protocol, a training sample containing 100,000 labeled examples was generated.

A modification of U-Net: SegNet [88], adapted for processing one-dimensional signals, consisting of two parts - an encoder and a decoder - was chosen as the neural network model architecture. A section of the input signal was subjected to downsampling procedure with the extraction of a feature map at each level. Then upsampling was performed with concatenation of feature maps obtained at the first step.

To evaluate the quality of the model performance, cross-validation was performed on a partitioned training dataset. Segmentation success was evaluated by F1 metric, which represents the harmonic mean between precision and recall. The F1 metric value of 0.84 was obtained, which demonstrated the ability of the model to correctly identify ATs on EEs in sufficient volume for further statistical analysis of the experimental data.

## Results and discussion

Application of algorithm on the basis of DL for processing of experimental data obtained by MEA method allowed to reveal the characteristics of changes in MBA of the isolated rat heart under the influence of LC and ADR. The U-net-based neural network localized ATs on the EEs despite significant distortions under the influence of ADR (Fig. 4, A) and/or LC (Fig. 4, B) of amplitude, shape, and frequency of occurrence of the ATs recorded by MEA electrodes.

**Figure 4.**
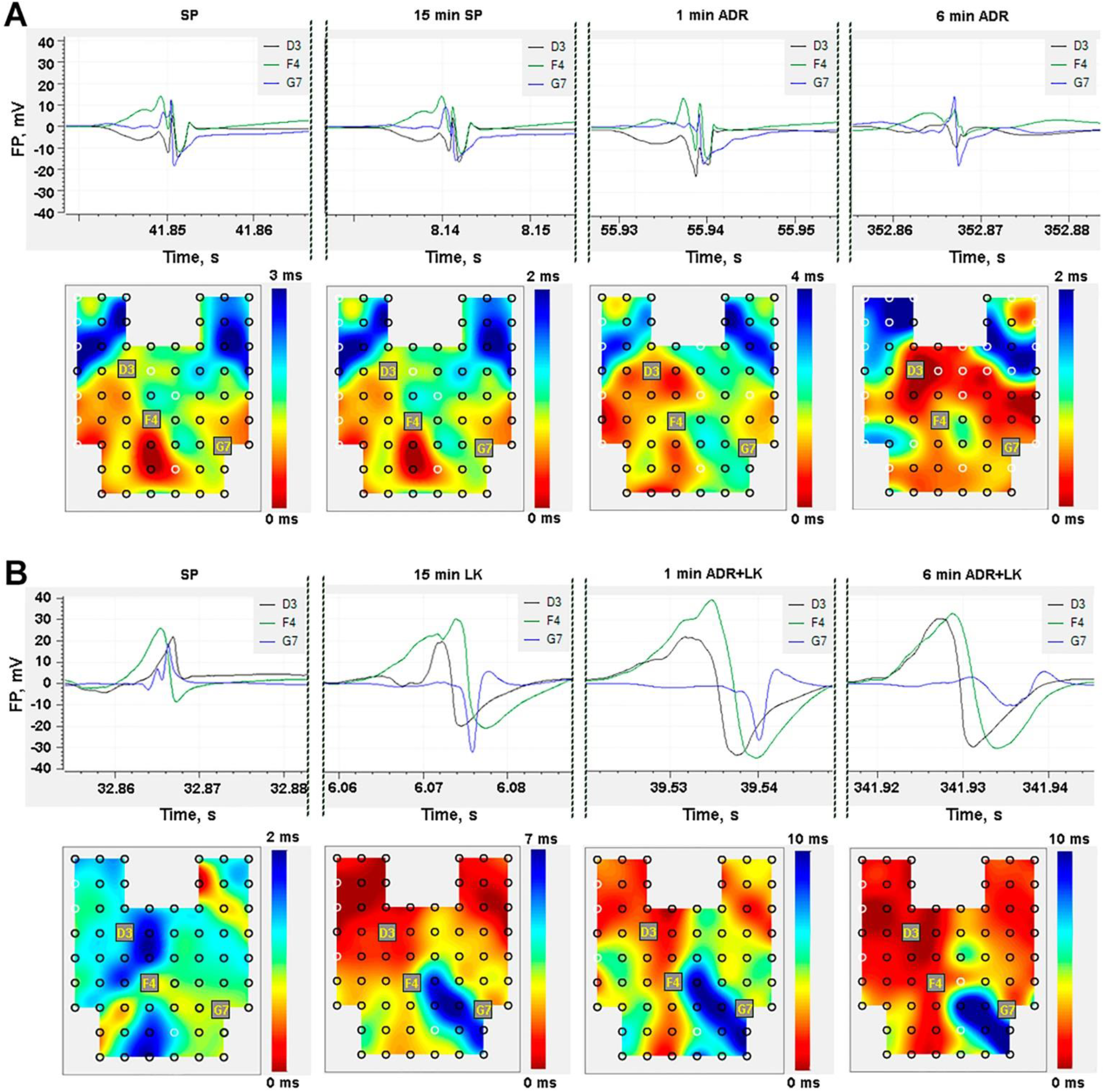
Changes of MBA of two isolated rat hearts during ADR exposure: A - in the «ADR» group, B - in the «LC» group; data are presented as EEs of three selected electrodes, the characteristics of LFPs on which change depending on the experimental conditions, activation maps showing changes in the pattern and rate of myocardial excitation spanning are also presented; black or white rings indicate MEA electrodes on which LFPs were or were not detected at the time of cardiac contraction, respectively (Cardio2D+, Multichannel Systems, Germany)

The performed experiments indicated that perfusion of the isolated rat hearts with a solution containing LC in the «LC» group for 15 minutes causes an increase in CCD and EPT, and a decrease in CBF. Meanwhile, under SP conditions in the «ADR» group, there were no statistically significant differences in these parameters between the 1st and 15th minutes of perfusion (p<0.05) (Fig. 5). The value of CV_CCD_ in both groups remained equal to 0.7±0.4%. At the beginning of the experiment for both groups were observed: CCD = 231±24 ms, EPT = 3.2±1.2 ms, CBF = 17.2±2.5 ml/min.

**Figure 5.**
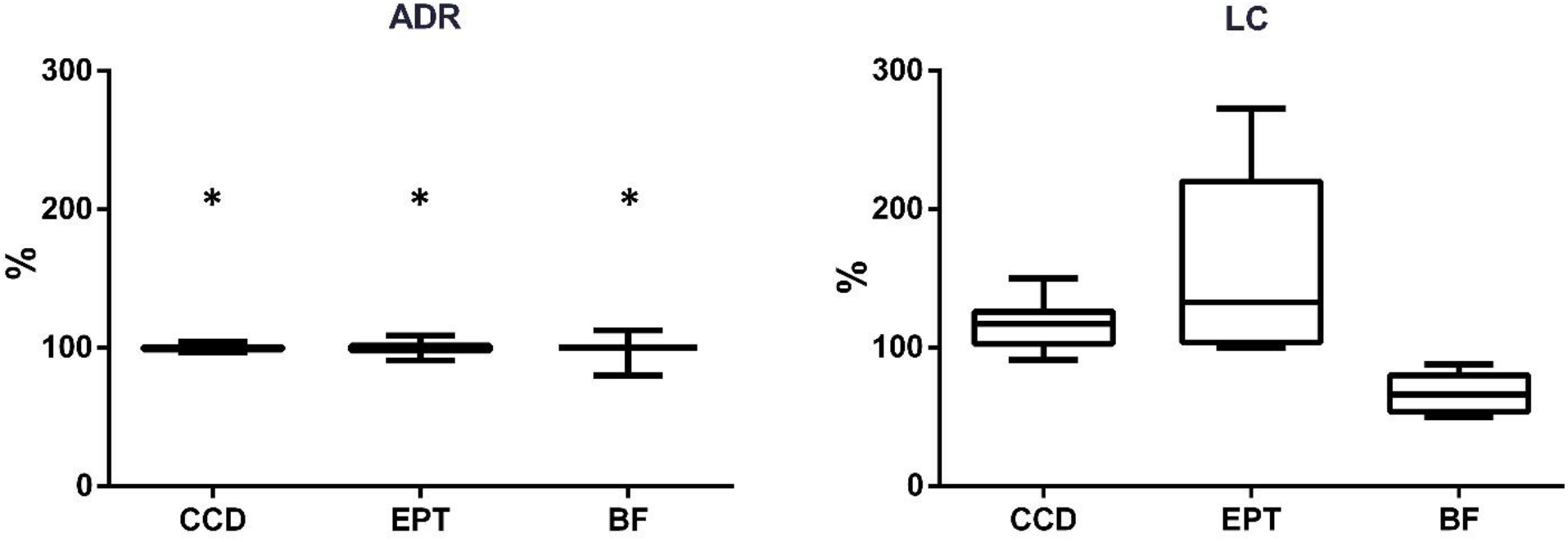
Percentage change from initial values of CCD, EPT and CBF parameters of the isolated rat hearts after 15 minutes of SP in the «ADR» group and by LC-containing perfusion solution in the «LC» group before ADR administration («whiskers» - range of variation, boxplot boundaries - interquartile range divided by median); * - no statistically significant differences by Wilcoxon’s criterion (p<0.05)

The observed negative chronotropic effect of additional LC in the heart has been demonstrated previously [89, 90]. In our experiments, we found an increase in LFP duration and amplitude after 15 minutes of LC perfusion (Fig. 4). From one point of view, we can assume that this phenomenon is advantageous in contrast to the decrease in action potential amplitude and duration, which may be caused by the suppressive effect of long-chain acylcarnitines on MBA [91]. In case of excessive accumulation, these products of fatty acid degradation, the concentration of which is controlled by LC, become the cause of cardiac dysfunction [64, 92, 93]. From another point of view, LFP alterations may be a sign of MBA abnormalities caused by decreased CBF.

According to literature reviews [67, 94], supplemental doses of LC and its derivatives (propionyl-LC, LC-L-tartrate) contribute to the improvement of cardiac function and on CBF, especially in pathologic conditions. The ability of supplemental LC to stimulate postischemic blood flow recovery and revascularization and to enhance endothelial function has been experimentally established [71, 73, 95]. The results of a number of studies confirm the vasodilating properties of LC [96-99]. However, no information on the use of LC as a vasodilator could be found in the scientific literature.

The concept has been proposed that the regenerative effect of LC in muscle tissues is related to its ability to stimulate blood supply through improved epithelial function [70, 72]. In further investigation of this concept, Spiering B.A. et al. [100] revealed a decrease in muscle oxygenation in athletes taking LC, which was attributed to increased oxygen consumption by muscle tissue. Thus, the vasodilatory effect of LC may not be sufficient to supply the oxygen demand of the tissue. The raised oxygen consumption upon exposure to supplemental LC may be related to the activation of aerobic respiration due to both intensification of fatty acid oxidation and enhanced glycolysis. Since LC is able to lower the acetyl-coenzyme A (CoA)/CoA ratio, thereby maintaining the activity of the pyruvate dehydrogenase complex and the oxidation of pyruvate formed from glucose [66, 93, 101].

Additional doses of LC have been shown to increase the strength of cardiac contractions [89, 102], and this may cause a decrease in CBF, which depends on the pressure on the vessels from the myocardium [103].

Thus, the slowing of excitation propagation in the myocardium detected in the present study may be caused by oxygen deficiency in experimental conditions during intensification of aerobic respiration and reduction of CBF caused by the addition of LC to the perfusion solution.

It is also conceivable that the increase in cardiac contractile force could have been due to an increase in Ca^2+^ ion concentration, which contributed to a decrease in electrical conduction velocity by blocking gap junctions [104, 105]. But in the works of Gökçe Y. [102] simultaneously with the positive inotropic effect of prolonged supplemental LC administration was demonstrated acceleration of calcium dynamics, preventing excessive accumulation of Ca^2+^ ions in cardiomyocytes.

In the experiments performed, a LC concentration of 1 mMol was used for perfusion of the isolated rat hearts. The experimental protocols of other authors suggested the use of LC in concentrations of 0.5-10 mMol [66, 106]. Based on literature data, the LC content in rat plasma and myocardium is comparable to that in humans and amounts to 45 μMol/L and 1-4 μMol/g wet weight, respectively [66, 107, 108]. Since LC transport into cardiomyocytes from plasma is limited by the work of protein transporters [50, 51], the chosen concentration should not have been excessive. In a preliminary series of experiments, the use of perfusion solution with lower LC content had no effect on the performance of the isolated rat heart. From literature reviews devoted to LC, it is known about the efficacy of long-term supplementation of this compound [81, 109]. Nevertheless, in the study by Broderick T.L. et al. [66], 60-minute perfusion of the isolated rat hearts with LC solution at a concentration of 10 mMol resulted in a more than twofold increase in the LC content in the myocardium [66], which confirms the probability of LC action in our experiments.

Administration of ADR under SP conditions induced in the isolated rat hearts the previously described effects of increased instability and HR (decreased CCD) [110]. In hearts perfused with LC solution after ADR administration, CCD values remained at a higher level – HR increased less intensively (Fig. 6) and also occurred more unstable than in the «ADR» group. The CV_CCD_ reached 16.8% in the «ADR» group and 18.5% in the «LC» group. Thus, the presence of LC in the perfusion solution did not lead to HR stabilization, as has been shown in studies by other authors when studying the action of LC in arrhythmias [111, 112].

**Figure 6.**
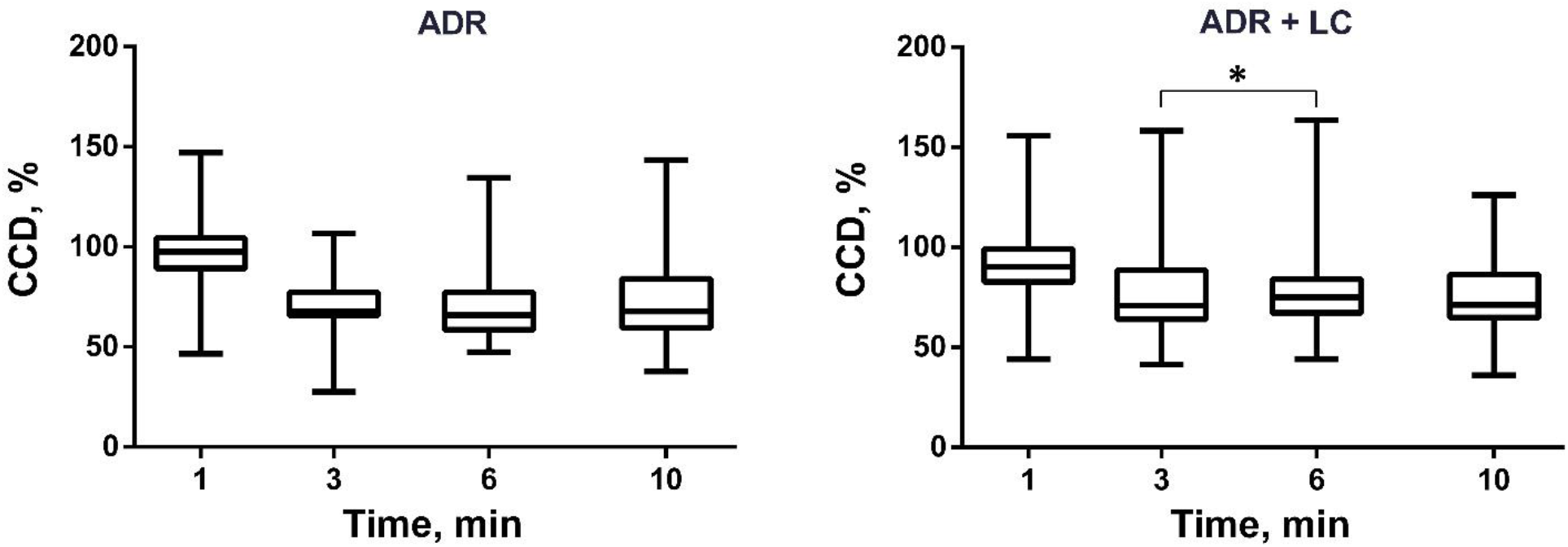
Dynamics of CCD changes of the isolated rat hearts under ADR exposure in «ADR» and «LC» groups as a percentage of the values of the parameter before ADR administration («whiskers» - range of variation, boxplot boundaries - interquartile range divided by median); * - absence of statistically significant differences according to the results of two-way ANOVA and Tukey’s test (p<0.05)

In the «ADR» group, administration of ADR in many hearts caused an episodic increase in EPT (slowing of conduction velocity) (Fig. 7). In the «LC» group, EPT values remained closer to the previous level after ADR administration as compared to the «ADR» group (Fig. 7), but these EPT values were already elevated (Fig. 5). The findings of the investigations show the antioxidant effect of LC [61], which could prevent the negative effects of ADR oxidative metabolites on MBA [113]. This may explain the maintenance of myocardial electrical conductivity at a stable level in the «LC» group when exposed to ADR.

**Figure 7.**
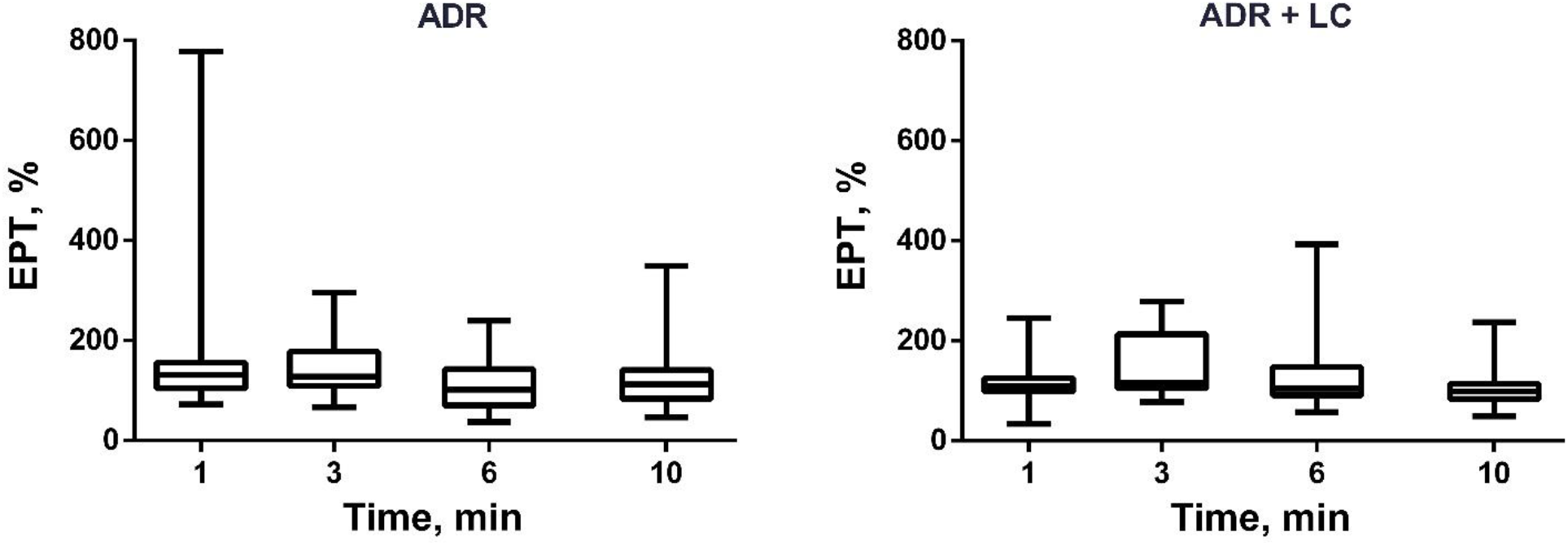
Dynamics of EPT changes of the isolated rat hearts under ADR exposure in «ADR» and «LC» groups as a percentage of the values of the parameter before ADR administration («whiskers» - range of variation, boxplot boundaries - interquartile range divided by median); the differences are statistically significant according to the results of two-way ANOVA and the Tukey test (p<0.05)

In both groups, ADR administration caused a decrease in CBF in the first minute, more significant in the «ADR» group in the absence of LC. An increase in CBF was then observed in the «ADR» group (Fig. 8). Initial CBF changes in the «ADR» group could be provoked by α-adrenoreceptor activation, which caused vascular smooth muscle contraction. The subsequent activation of β-adrenoreceptors caused vasodilation [114]. No statistically significant (p<0.05) increase in CBF was found in the «LC» group. Hence, in the «LC» group, there was not only a decrease in CBF under SP conditions, but also a suppression of vascular reactivity that should be observed in response to ADR action, as occurred in the ADR group.

**Figure 8.**
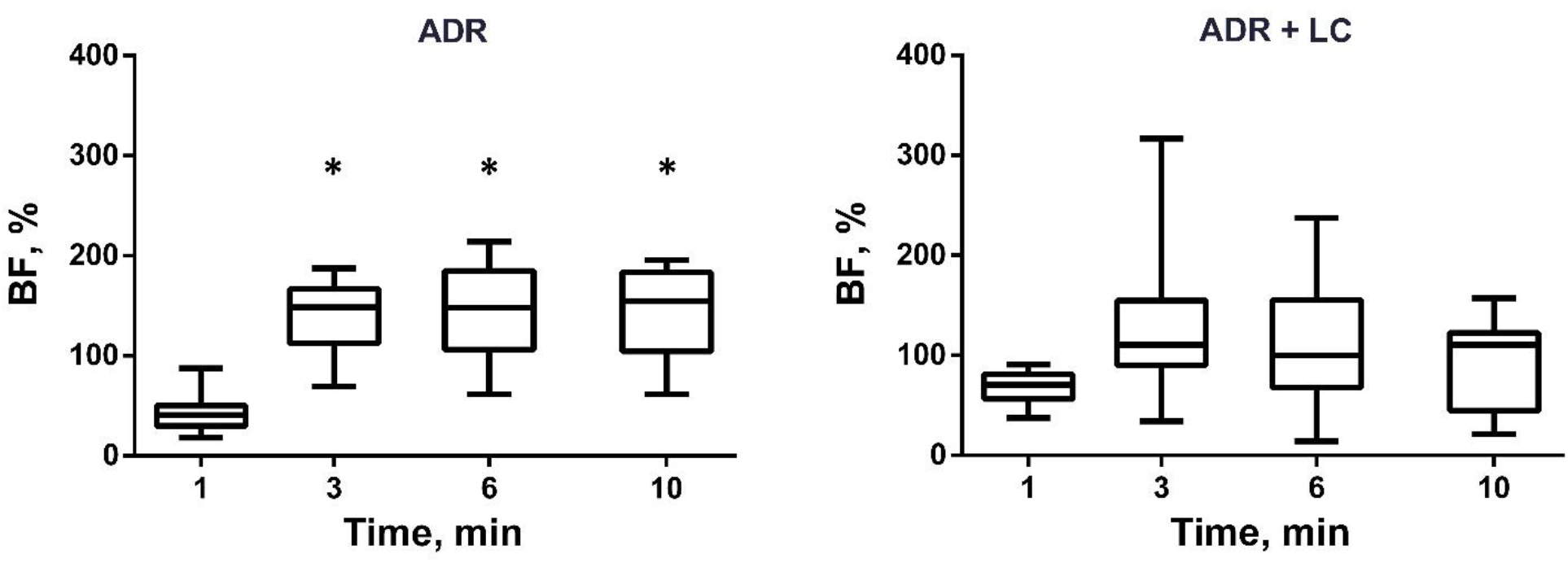
Dynamics of CBF changes of the isolated rat hearts under ADR exposure in «ADR» and «LC» groups as a percentage of the values of the parameter before ADR administration («whiskers» - range of variation, boxplot boundaries - interquartile range divided by median); * - absence of statistically significant differences according to Friedman’s test and Dunn’s test (p<0.05)

**In conclusions**, the application of the neural network model based on U-Net architecture provided an efficient analysis of the experimental data obtained by myocardial MEA mapping and new findings about the effects of additional doses of LC on cardiac function. In particular, new observations of the influence of LC on the spatio-temporal characteristics of MBA of the isolated rat heart have been demonstrated.

The results of the present studies showed that acute administration of additional doses of LC can cause a decrease in HR and excitation conduction velocity in the myocardium, as well as in the intensity of coronary supply. These effects had no significant suppressive effect on cardiac function *ex vivo*, but prevented a proper response to ADR action, which should be manifested by increased HR and enhanced coronary circulation.

According to these findings, it can be concluded that in the presence of CVD and under conditions of intense physical activity, the dose and administration of LC-based drugs should be selected with caution, as the cardiovascular system may not be prepared for the ADR release induced by emotional or physical stress under the influence of these compounds.

Thus, this study demonstrated how a neural network model based on the U-Net architecture can be used to automate the process of localizing ATs on multiple MBA electrograms recorded with MEA from the epicardium of a contracting isolated rat heart perfused by the Langendorff method.

